# A multi-step completion process model of cell plasticity

**DOI:** 10.1101/2024.10.01.616127

**Authors:** Chen M. Chen, Rosemary Yu

## Abstract

Plasticity is the potential for cells or cell populations to change their phenotypes and behaviours in response to internal or external cues. A common way to study a plasticity programme is to track the underlying molecular changes over time, by collecting omics time-series data. However, there remains a lack of mathematical models to elucidate and predict molecular behaviours in a plasticity programme using omics time-series data. Here we report a new mathematical framework that models cell plasticity as a multi-step completion process, where the system moves from the initial state along a path guided by multiple intermediate attractors until the final state (i.e. a new homeostasis) is reached. In benchmarking tests, we show that our developed Control Point (CP) model fits omics time-series data and identifies attractor states that are well-aligned with prior knowledge. Importantly, we show that the CP model can make non-trivial predictions such as the molecular outcomes of blocking a plasticity programme from reaching completion, in a quantitative and time-resolved manner.

**Significance statement:** Cell plasticity is fundamental to a wide range of complex biological processes, for example in embryonic development, immunology, regenerative medicine, aging, cancer, and others. We developed a new mathematical model that leverages omics time-series data to understand and make accurate predictions about cell plasticity. This can lead to new discoveries and open up new avenues of research in medicine and bioengineering.

## Introduction

Cells and cell populations are highly plastic, meaning that they can robustly respond to a variety of signals and (non-lethal) perturbations by changing their characteristics and behaviours ^1,2^. These phenotypic changes arise through a cascade of molecular events over time, and is ‘complete’ when a new homeostasis is reached ^2,3^. To study plasticity, it is common to systematically track the time course of these molecular changes by collecting omics (e.g. transcriptomics, proteomics) time-series data. However, mathematical models of plasticity using omics time-series data are lacking. Typical methods to extract information from omics time-series data, such as time-lagged correlation ^4^, molecular event inference ^5^, and interaction/regulatory network reconstruction ^6^, are largely descriptive and have limited use in making predictions that are quantitative and/or time-resolved.

Many molecular programmes related to plasticity can be viewed as a multi-step completion process: the system moves, over time, from the initial state along a path guided by multiple intermediate attractors, until the final state is reached ^7,8^ (Fig 1A-C). Here we describe a new mathematical framework that models the molecular changes occurring during such a completion process, i.e. omics time-series data, as a function of the stochastic probabilities of reaching each attractor state over time. We show that our developed Control Point (CP) model accurately identifies attractor states by their molecular markers and timing. More importantly, the CP model can be used to make non-trivial predictions about the modeled plasticity programme. For example, to date no modeling framework is able to predict the molecular outcomes of a plasticity process if it is blocked from reaching completion. We show that the CP model can make such predictions in a quantitative and time-resolved manner, to an R^2^ of 0.53-0.63.

**Figure 1.**
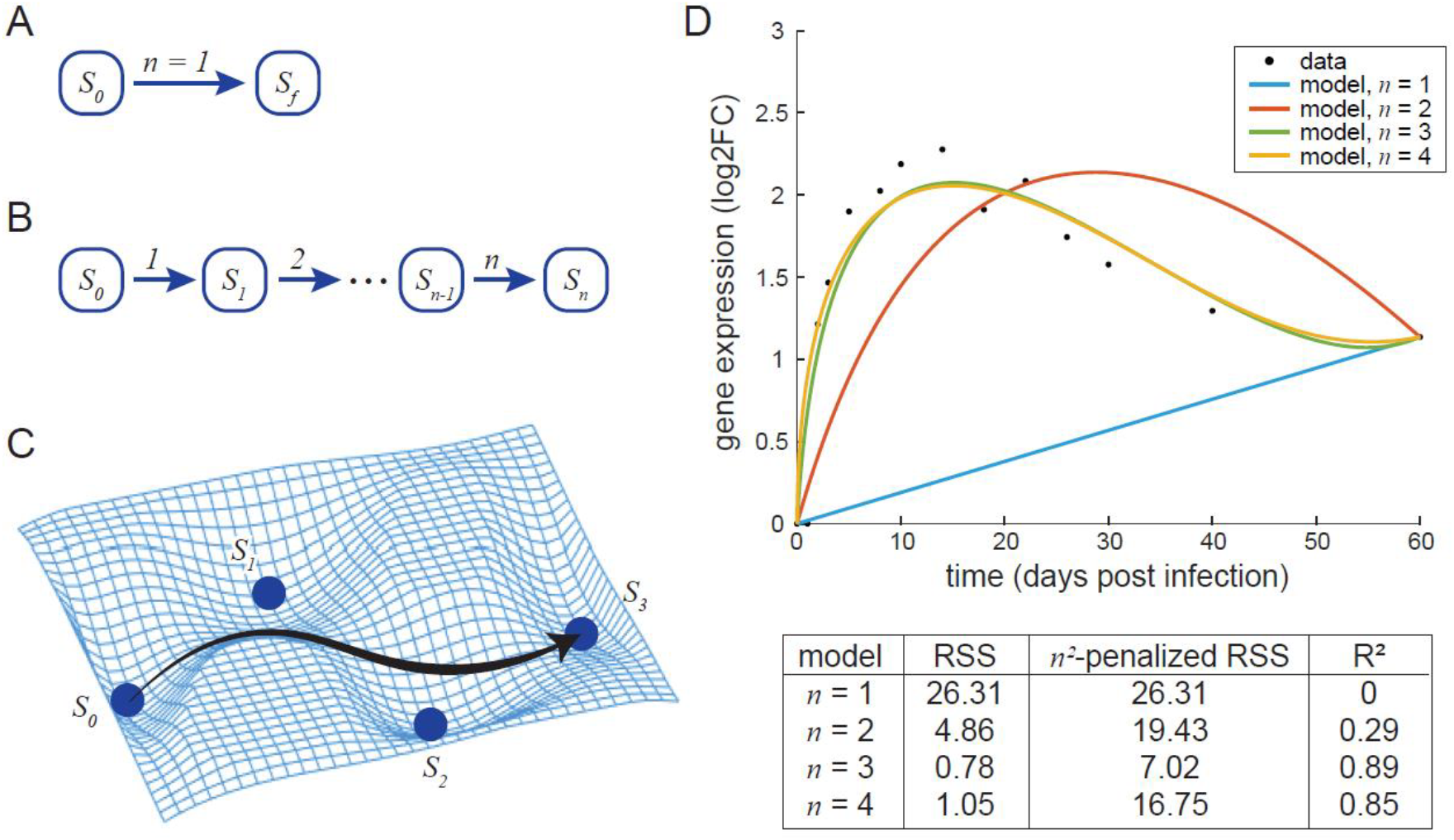
Modeling cell plasticity as multi-step completion processes. (A), schematic of the simplest completion process, where the system reaches the final state in single step (*n* = 1). (B-C), schematic of an *n*-step completion process where the system moves along a path guided by *n* −1 intermediate attractors, until reaching the final state. (D), best fit models of *n* = 1, …, 4, of a single gene (*Il4i1*) from ref ^10^. Data points are expression levels of *Il4i1* in the mouse lung for 60 days following a viral infection (ref ^10^). Colored lines are fitted CP models for the single gene. The residual sum of squares (RSS), *n*^*2*^-penalized RSS, and R^2^ of each fitted model are given.

## Results

Consider some plasticity programme that can be represented by a completion process as shown in Fig 1A-C, where each attractor state *S* is ‘marked’ by a particular set of molecular characteristics. We model this process as *n* discrete state transitions, i.e. *n* independent Bernoulli trials, where a ‘success’ means that the system proceeds to a particular state, while a ‘fail’ means that the system stays in a previous state. At any moment during the progression of the process (τ ∈ [0,1]), the probability of successfully undergoing exactly *k* state transitions, with *k* = 0, …, *n*, is given by the binomial distribution ^9^ (see Methods section, eq. 1). Suppose that omics data (e.g. transcriptomics) is collected over time, at enough time points to be fully representative of the completion process. Then, for each molecular characteristic *g* measured over time (e.g. abundance of a transcript), we find a model of a size of *n* that fits the measurement data arbitrarily well (Fig 1D), and solve for a set of control points *P*_*g*_ that specifies this model (see Methods section, eq. 2). Then, for all measured molecules *G* ∋ *g*, we use the collective control points *P*_*G*_ to infer the intermediate attractor states (*S*_*k*_, *k* = 1, …, *n* − 1) by both their timing (*t*) and their molecular markers (a subset of the measured molecular characteristics *G*). Note that at *k* = 0 and *k* = *n*, attractors *S*_0_ and *S*_*n*_ represent the initial and final states of the process, respectively, and their markers are taken as all measured molecules *G*; see details in the Methods section. Below we illustrate these concepts, and benchmark the CP model, using two molecular programmes in plasticity as examples.

In the first example, we model a viral infection-induced gene expression response in the mouse lung ^10^, representing a plasticity programme at the tissue level. Response to a viral infection is a textbook example of a multi-step completion process, consisting of 4 phases between the initial state (*S*_0_) and final recovery to homeostasis (*S*_*n*_): first the 3 phases of the immune response (innate response, T cell response, and B cell response), followed by a 4^th^ phase of tissue repair ^11^. To model this process, we mined a set of transcriptomics time-series data of the mouse lung collected after a non-lethal infection with influenza A over a period of 60 days ^10^. We find the best-fit model for each gene *g* over time (see Fig 1D for an example), and use a kernel density plot for a global overview of the positioning of the control points *P*_*G*_ of all genes along the time-axis (Fig 2A). The CP model accurately captures that this plasticity programme is best described by a completion process of 4 intermediate attractor states between the initial state *S*_0_ and the final state *S*_*n*_ (Fig 2A), in agreement with the known 4 phases of the viral infection-induced response. Moreover, the model-identified molecular markers of each attractor state are fully in agreement with known markers of the different phases of the viral infection-induced response (Fig 2A). This includes *Ifi44* and *Ift1* as markers of the first attractor state (innate response)^12,13^; *Cd8a* and *8b1*, classic markers of cytotoxic T cells ^11^, marking the second attractor state; the B cell marker *Cd79b* ^14^ and 14 out of 16 detected *Igh, Igj*, and *Igk* genes indicative of antibody production during the B cell response ^11^, marking the third attractor state; and *Six1, Bcas2*, and *Dmbt1* marking the final attractor state of tissue repair ^15-17^. Finally, the timing of each phase of the response, modeled in a completely data-driven manner (Fig 2A), perfectly recapitulates schematics of the host immune response over time that can be found in textbooks ^11^.

**Figure 2.**
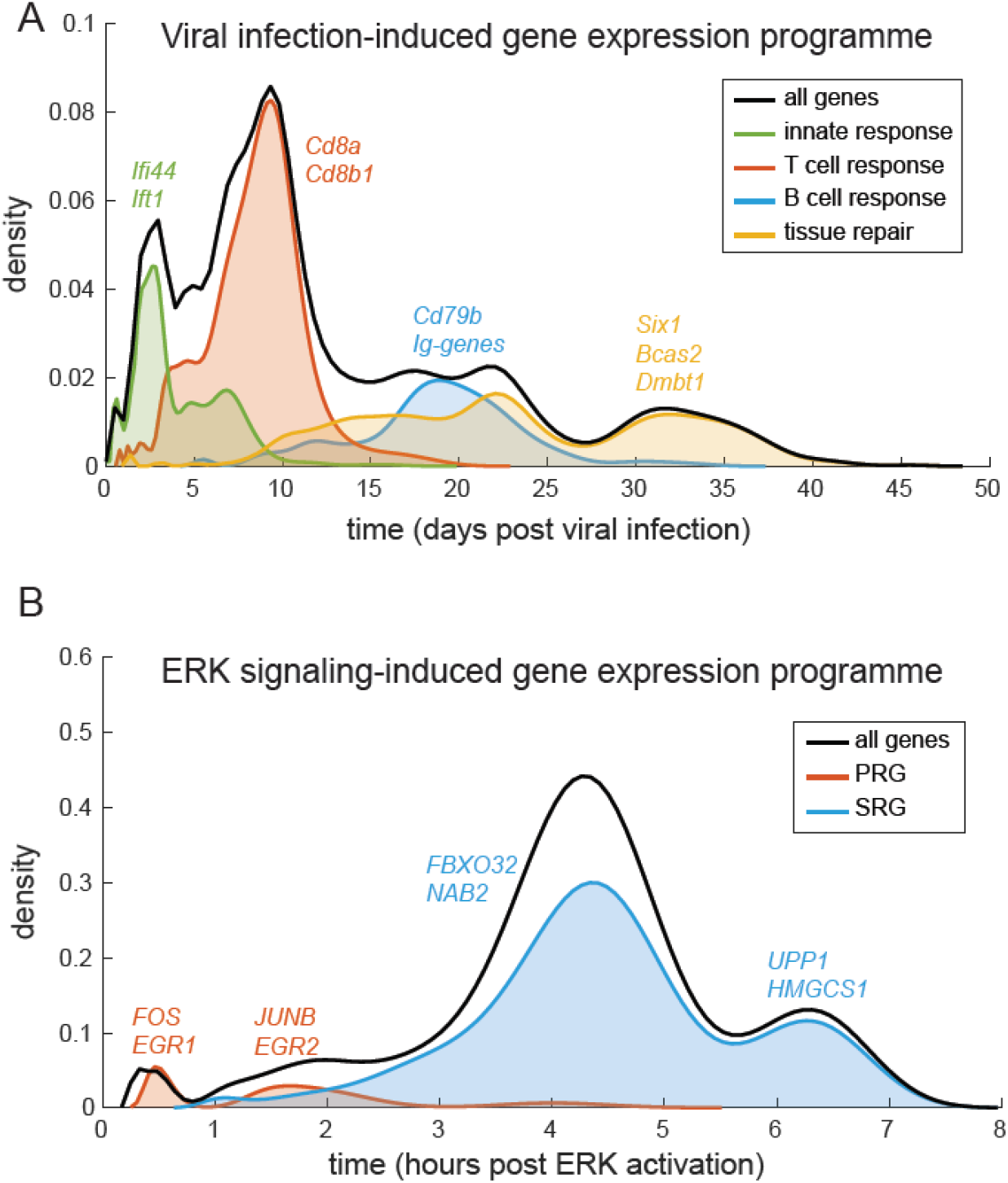
The CP model accurately recapitulates known phases of plasticity programmes, including their timing and molecular markers. (A), CP model-identified attractor states of a viral infection-induced response in the mouse lung, from ref ^10^. (B), CP model-identified attractor states of ERK signaling-induced gene expression in HEK293 cells, from ref ^18^.

In the second example, we use the CP model to study ERK signaling-induced gene expression in HEK293 cells ^18^. ERK is the final kinase in the classic RAF-MEK-ERK signaling pathway. Upon activation of ERK signaling, transcriptomic changes of HEK293 cells (and other cell lines) occur through two phases: first the induction of a set of genes called primary response genes (PRGs), followed by the induction of secondary response genes (SRGs) which is dependent on the translation of the PRGs ^18,19^. To model this programme, we mined a set of ERK signaling-driven transcriptomics time-series data in HEK293 cells, covering a span of 10 hours ^18^. The CP model identifies two attractor states marked by PRGs, followed by two attractor states marked by SRGs, recapitulating the known phases of ERK signaling (Fig 2B). Interestingly, this analysis shows that the SRGs, which are previously thought of as a single group of genes characterized by their dependence on PRG translation, are in fact marking two temporally distinct attractor states (Fig 2B). The earlier of the two SRG-marked attractor states, at 3.5-5 h after ERK signal induction, is marked by genes such as *FBXO32* and *NAB2*, which are negative regulators of ERK signaling and/or its downstream effectors ^20-22^. The later attractor state, at 6-7 h after ERK signal induction, is marked by genes such as *UPP1* and *HMGCS1*, which are metabolic enzymes supporting cell proliferation ^23,24^. The discovery of these temporally and functionally distinct attractor states therefore suggests a division of the SRGs into two subsets for further molecular characterization – similar to what has been done in the investigation of the PRGs ^18,19^.

Previously methods to identify the phases of ERK signaling-induced gene expression (e.g. peak expression time ^19^) are not able to make non-trivial predictions. Here we show that the CP model can quantitatively predict the molecular outcome of the modeled completion process, if the process is blocked from reaching completion. Conceptually speaking, blocking the completion process at step *k* (i.e. blocking the transition from *S*_*k*−1_ to *S*_*k*_) would predict that the system becomes attracted to the preceding state, *S*_*k*−1_, as the new stable attractor 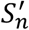 (see Fig 1B-C). Using the CP model, the dynamics of the resulting molecular programme can be predicted in a time-resolved manner. To illustrate this, we used a published dataset where HEK293 cells are co-treated with an ERK activation signal and a protein synthesis inhibitor for 1, 2, and 4 hours ^18^. The protein synthesis inhibitor blocks the translation of PRGs into proteins, thereby preventing the induction of SRGs which marks the next attractor state(s) ^18^. Using these experimental gene expression measurements as validation, we demonstrate that the CP model predictions are quantitative, time-resolved, and accurate to an R^2^ of 0.53-0.63, a dramatic improvement to the null model which is completely unable to make such predictions (R^2^ = 0, Fig 3).

**Figure 3.**
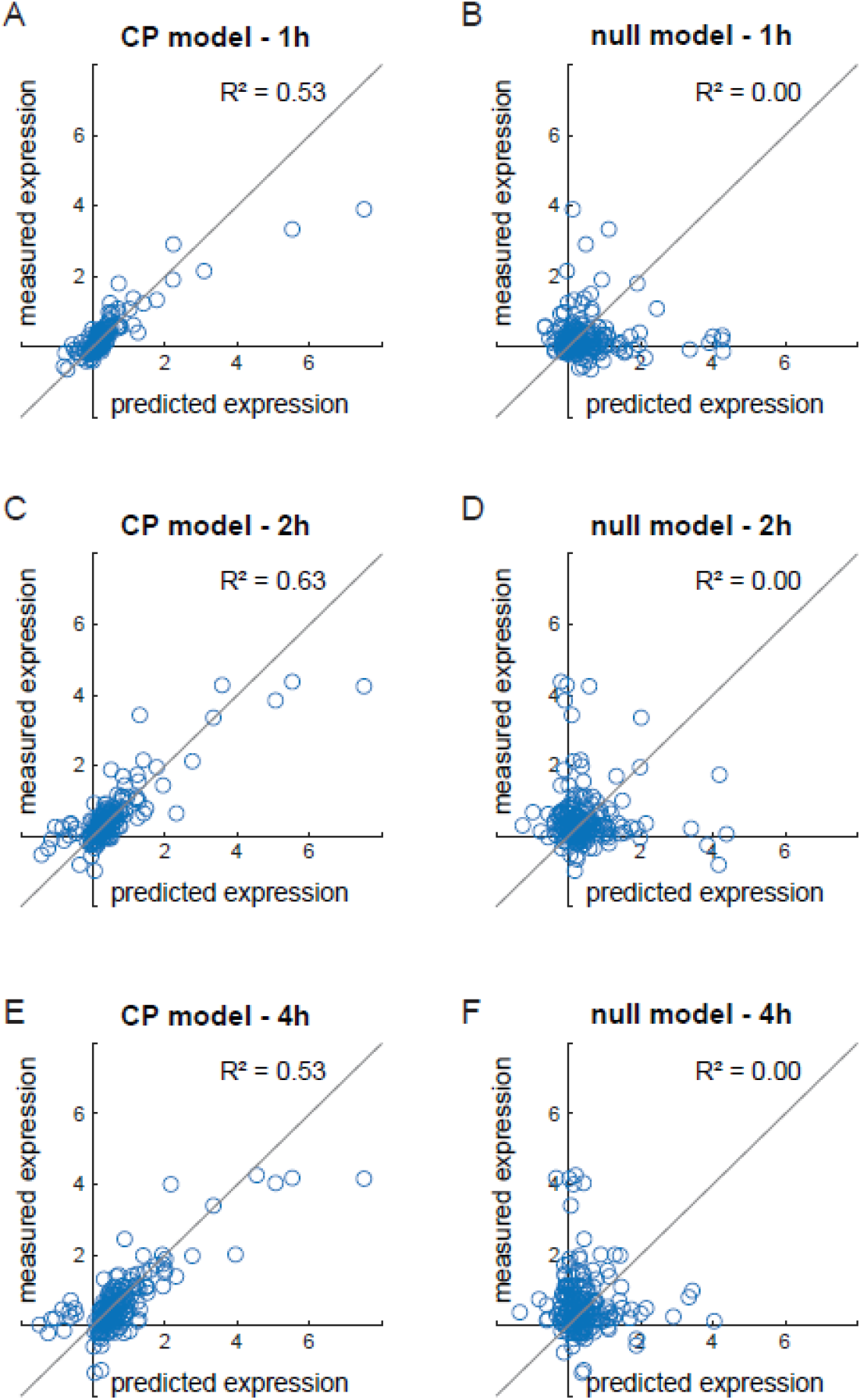
Non-trivial predictions of the CP model. (A, C, D), comparison between model-predicted and experimentally collected gene expression in the ERK signaling-induced gene expression programme, when cells are treated with the protein translation inhibitor CYHX. The experimentally collected data are mined from ref ^18^. (B, D, F), comparison between a null model and experimental data mined from ref ^18^.

## Discussion

Here we describe a new modeling framework for cell plasticity, that models the molecular changes over time in a plasticity programme as a multi-step completion process. Using omics time-series data as input, the CP model accurately identifies the attractor states of a complex plasticity programme by their timing and molecular markers, in a completely data-driven manner. We benchmark the CP model using two molecular programmes in plasticity, showing that the model fits data well and identifies attractor states that are well-aligned with prior knowledge. Moreover, CP model-based analysis can lead to new knowledge discovery, for example showing that SRGs in the ERK signaling-induced gene expression programme can be divided into two subsets, marking two attractor states that are functionally and temporally distinct.

The CP model is the first description of a generalized modeling framework for cell plasticity that can make non-trivial predictions in a quantitative and time-resolved manner. We believe that this is the most important property of a useful model: the ability to not only accurately describe the molecular behaviours of a cell, but also to predict and eventually use these predictions in bioengineering or biomedical applications. Here we demonstrate that the CP model can predict the molecular outcomes of blocking a completion process from reaching completion. This can, for example, help direct the search for strategies to combat drug resistance acquisition in cancer cells (by modeling drug resistance acquisition as a completion process), which is a Cancer Grand Challenge that remains to be addressed ^25^.

A current limitation of the CP model is the assumption that the modeled process has a single path to completion (Fig 1A-C). For some plasticity programmes, this assumption may not hold true. For example, there is considerable redundancy in the growth factor-activated signal transduction (protein phosphorylation) network of a eukaryotic cell ^26^. Thus, a phospho-proteomics time-series dataset mapping the signal transduction of cells upon growth factor exposure, may be better modeled by a multi-path completion process ^8^. Future development of such models could rapidly advance our ability to understand and manipulate cellular signal transduction in diverse contexts.

## Methods

### Model description

Consider an *n*-step, single-path completion process as shown in Fig 1A-C. We model the *n* state transitions as *n* independent Bernoulli trials, where a ‘success’ means that the system proceeds to a particular state, while a ‘fail’ means that the system stays in a previous state. At any moment in the progression of the process, τ ∈ [0,1], the probability of successfully undergoing exactly *k* state transitions, with *k* = 0, …, *n*, is given by the binomial distribution ^9^, *B*_*n*,*k*_(τ):

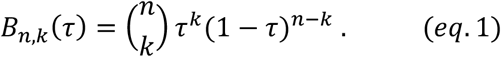

The mathematical expression of the entire *n*-step completion process is:

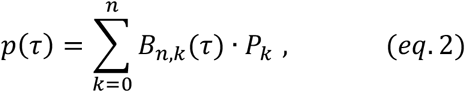

where *p*(τ) denotes the molecular characteristics (omics) of the system over time (to be fitted), and *P*_*k*_ denotes the (fitted) *k*^th^ control point of the molecular characteristics. Note that B_*n*,*k*_(τ) is a complete basis (see Bernstein’s proof of the Weierstrass approximation theorem, ref ^27,28^), thus every dataset can be fitted arbitrarily well by some (possibly very complex) CP model. Additionally, *p*(τ) is uniquely determined by its control points *P*_*k*_: if two functions *p*(τ) and *p*′(τ) are identical, then 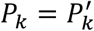 by definition ^29^. In practice, we fit the CP model to each molecular characteristic *g* independently (i.e. solving for each *g* a set of control points {*P*_*g*_} = *P*_*g*,*k*_, *k* = 0, …, *n*), since not all characteristics are expected to be a marker for every intermediate state. The intermediate states of the entire completion process are then inferred through collective analysis of the characteristic-specific control points, described below.

### Input data and pre-processing

#### Data selection and transformation

The input data can in principle be time-series data of any type of molecular characteristics (i.e. omics), sampled at enough time points to be fully representative of the completion process. To avoid overfitting, we recommend at minimum 3*m* + 2 data points, where *m* = *n* − 2 is the estimated number of intermediate states based on prior knowledge. Taking transcriptomics as an example, since transcripts that are not differentially expressed over time are not of interest (they would not be markers of any intermediate state), data selection is performed by standard differential expression analysis appropriate to the data type and process of interest. For datasets that contain more experimental noise, additional data filtering can be performed. For example, for the viral infection-induced immune response dataset ^10^, we further filter out genes with < 5 measurement timepoints (out of 14 total) with a significant differential expression. All data are transformed to log2 fold change against the unperturbed control sample at *t*_0_.

#### Identifying time of final expression level

For time-series omics datasets of plasticity, we expect conservative experimental procedures where more data is collected over a (slightly) longer timeframe than the completion process itself. Moreover, for each molecular characteristic *g* measured in the omics timeseries dataset, we consider the possibility that *g* reaches its final expression level before the entire process is complete. Therefore, for each *g*, we determine the time at which final expression is reached, by evaluating each measurement *y*_*g*_ against the range of *y*_*g*,*final*_ ± *φ* · (*y*_*g*,*max*_ − *y*_*g*,*min*_), with an arbitrarily small *φ* ∈ (0,1). In reverse temporal order, we find a set of timepoints where all consecutive *y*_*g*_ is within the range as given above. Then, within this set, the *j*^*th*^ timepoint (*j* being an arbitrary small integer, default 2) is taken as the time of final expression, *t*_*final*_.

#### Calculation of τ

τ is a scaled ‘progress’ parameter that ranges from 0 to 1. For each *g* with some measured values (or bootstrapped values, see below) over time, as a set of points in the vector space of time and the molecular measurement *p*_*g*,*i*_ = (*t*_*i*_, *y*_*g*,*i*_), first we take the scaled distance between each consecutive datapoint, *d*(*p*_*g*,*i*_, *p*_*g*,*i*+1_) · λ. The scaling factor λ is introduced to control the different scales between *t* (e.g., tens to hundreds of days or hours) and *y* (as a log2 fold change, typically within ± 10). We then calculate the total distance *D* = ∑_*i*_ *d*, followed by calculating 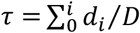, such that τ = 0 at *t*_0_, and τ = 1 at *t*_*final*_.

#### Data bootstrapping

Weighted bootstrap is performed for each *g* to generate evenly spaced datapoints prior to model fitting. We bootstrap data to be evenly spaced on τ, i.e. consecutive (bootstrapped) datapoints have equal distances. For each bootstrapped datapoint, a default 1,000 random samples is taken from all *y*_*g*_, with measurements taken at a closer time to the bootstrap timepoint receiving higher weights. The average of the 1,000 (weighted) random samples is then taken as the bootstrapped datapoint. By default, 100 datapoints are bootstrapped.

### Model fitting and analysis

#### Model fitting

We use multiple linear regression with the binomial distribution (eq. 1) as the basis (also known as the Bernstein basis, see ref ^27,28^) to fit CP models with increasing *n* to the (bootstrapped) data for each *g*. Of note, we take the measurements of *g* at *t*_0_ and *t*_*final*_ as fixed control points, thus for *n* = 1 the model is not fitted but directly evaluated by the linear correlation between the data and the model. If this linear correlation is poor (by an arbitrary cutoff), then models for *n* > 1 are fitted by multiple linear regression, using the built-in MATLAB function `regress`, which outputs (*n* + 1) number of control points for the model *P*_*k*_, *k* = 0, …, *n* (see eq. 2). The maximum *n* allowed is the number of timepoints with *measured* data for *g*, although in practice, we further reduce maximum *n* by a factor of 2 or 3 to save computational time, since models with large *n* are almost definitely overfitted. For each fitted model, goodness-of-fit parameters are then calculated, with the *measured* values of *g* only, including the residual sum of squares (RSS); RSS penalized by a factor of *n*^2^; and the coefficient of determination (R^2^).

#### Model selection

Based on the goodness-of-fit parameters from model fitting, the model that best fit the data can be selected by either minimum *n*^2^-penalized RSS, or an arbitrary cutoff of RSS or R^2^. We find an arbitrary cutoff of R^2^ to be the most intuitive selection criteria, and facilitates the comparison between all *g* within a dataset or even between datasets.

#### Global analysis and visualization

Following model fitting and selection, for each measured molecule *g* we have a set of optimal control points to describe the changes in *g* over time, {*P*_*g*_}. For all measured molecules in the omics dataset, then, we have {*P*_*G*_ }, *G* ∋ *g*. We define attractor states *S* by the relative density of {*P*_*G*_ } along the time-axis, where subsets of *P*_*G*_ cluster more densely together on the time-axis. The clustering of *P*_*G*_ on the time-axis infers both the global number of attractors as well as their timing during the completion process, while the ‘position’ of ⟨*P*_*G*_ ⟩ along the multi-dimensional ‘axes’ of omics data represents the molecular markers (subset of all measured molecules *G*) of each attractor *S*. This is visualized by a kernel density plot on the time-axis. Here, the timing and molecular markers of each model-identified attractor *S* is then compared to the known markers of each phase of the plasticity programmes, showing excellent agreement between the CP model and prior biologically knowledge about these programmes. In an unknown or understudied plasticity programme, the CP model-identified attractor states would represent new knowledge discovery, which can in turn be validated and subject to further study.

#### Model predictions

To predict the molecular outcome of interrupting the completion process at step *k* (i.e. blocking the transition from *S*_*k*−1_ to *S*_*k*_), each molecular characteristic *g* is evaluated by the subset of its control points *P*_*g*,*i*_, where *t*_*i*_ < *t*_*k*_. By eq. 1, this gives the dynamics of the interrupted process as predicted values in the vector space (*t, y*_*g*_), i.e. the predicted expression values of *g* over time *t*, for τ ∈ [0,1]. To make exact predictions of the expression value of *g* at a precise moment *t*, we first evaluate eq. 1 in the *t*-axis to solve for the corresponding τ, then calculate the expression value of *g* by evaluating eq. 1 in the *y*-axis. The coefficient of determination R^2^ is calculated to compare the predicted gene expression values by the experimentally measured values ^18^. As comparison, a null model is constructed by a random permutation of the experimentally measured gene expression values in the CYHX-dataset ^18^. R^2^ values between the null model predictions and the experimental values are calculated to be negative, indicating that the predictive power of the null model is very poor (residual sum of squares > total sum of squares). For ease of interpretation, we take these negative R^2^ values as R^2^ = 0.

## Data and code availability

The CP model is available as a suite of MATLAB functions at https://github.com/Radboud-YuLab/CPmodel. All mined data used in this paper, and scripts for data visualization, are provided at https://github.com/Radboud-YuLab/CPmodel. All procedures are implemented in MATLAB R2021a.

## Acknowledgements

The Yu lab is supported by grants from the Dutch Cancer Society and the Radboud-Western Collaboration Fund.

